# Comment on “Solving the time-dependent protein distributions for autoregulated bursty gene expression using spectral decomposition” [J. Chem. Phys. 160, 074105 (2024)]

**DOI:** 10.1101/2025.02.05.635946

**Authors:** Alexandre F. Ramos

## Abstract

The authors of the commented article claim that the exact time-dependent solutions of the stochastic model for a self-repressing gene previously obtained by us are both “incomplete and incorrect” because their calculations issued complex numbers as decaying rates of the system to steady state. We show that the imaginary decaying rates result from a methodological artifact. We use a linear algebraic approach to show that the decaying rates are those reported in [‘Exact time-dependent solutions for a self-regulating gene.’ Phys. Rev. E 83: 062902 (2011)]. Thus, our solution is both complete and correct. Additionally, the authors claim that they have discovered a new class of operator having complex eigenvalues and non-orthogonal eigenfunctions. We show that the operator can actually be written in the usual self-adjoint form, and, hence, it has real eigenvalues and orthogonal eigenfunctions.

## I. INTRODUCTION

In the commented article the exact solution of the model for the self-repressing gene[1] is classified as incomplete and incorrect : the authors found complex-valued decaying rates, and we, real-valued ones. They also raise doubts about the correctness of the symmetries underpinning the integrability of our model[2], but neglect their physical implications[3]. That is unexpected as symmetries are a main toolset of theoretical physics and for investigating the invariances[4] or degeneracies[5] of biological systems. Previously[6] a vigorous critique to the model presented in[7] was made while disregarding its phenomenological basis[8]. Because of an interpretative disagreement the authors (mis)classified as incorrect the exact solutions that we found[1, 7]. In our model, we consider a stochastic process having two random variables (*m, n*): *m* is the state of the promoter of the gene as ON or OFF, and *n* is the number of proteins in the citoplasm. The probability of finding the gene ON (or OFF) and *n* proteins is *α*_*n*_ (or *β*_*n*_). The time-dependencies of the probabilities are omitted. In[6] the authors proposed an alternative model for a self-repressing gene, and reported a closed expression for the steady state probabilities *α*_*n*_. However, no formulae for *β*_*n*_ or the normalization constant were obtained. Those missing functionals were calculated by us, and reported along with a discussion about the interpretation of stochastic models for gene regulation[9]. Now the authors combine the master equations for a bursty/non-bursty self-regulating gene[10] of which our model[7] is a particular case. Taking ansatz of separability and exponential time decay they obtain the time-dependent solutions. By truncating a continued fraction the authors found complex valued decaying rates contrasting with our real-valued ones[1]. Prima facie, this is contradictory with their calculations, and, again, the authors rushed in to criticize our results[1, 2].

In this comment we show that: the complex decaying rates result from a methodological artifact; our real-valued eigenvalues are correct; our solutions complete. The methodological artifact is shown using the stochastic model for the constitutive and binary externally regulated gene. A linear algebraic approach is employed as an alternative to demonstrate results in[1, 2].

## II. METHODS

Consider a system of first order coupled ODEs governing the dynamics of a column vector *Y* (*t*), 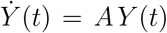 where *A* is a *n* × *n* matrix. *Y* (*t*) = *D* (*t*)*Y*_0_ is the solution of the system and *Y*_0_ is the initial condition. *D* (*t*) is the Dyson series, and, for *A* constant in time, *D* (*t*) = 1 + *At* + *A*^2^*t*^2^*/*2! + … = exp(*At*). This solution is unique and can be obtained numerically or analytically (Chap. 14[11]). The eigenvalues and eigenvectors of *A* may aid on computing exp(*At*) turning the solutions into eventually summable linear combinations of eigenvectors weighted by the exponentials of the eigenvalues. *Y* (*t*) reaches a steady state when *A* has real non-positive eigenvalues while damped oscillations occur for complex eigenvalues having negative real components.

## III. RESULTS

Case 1: a constitutive gene can be modeled as a birth and death stochastic process with rates *k* and *ρ*, respectively. The probability of nding *n* proteins at time *t, ϕ*_*n*_, obeys 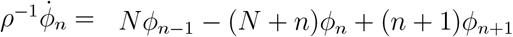 whose solution is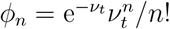, with *ν*_*t*_ = *N* + (*N*_0_ − *N*)e^−*ρt*^, *N* = *k/ρ*, and *N*_0_ the parameter of a Poisson process setting the initial condition. The generating function of *ϕ*_*n*_ is a linear combination of temporal-decaying exponentials, namely e^−*lρ t*^, *l* = 0, 1, …, *n*, that sums up as e 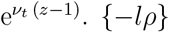 are eigenvalues of the transition matrix *U*_*c*_ having as entries the transition probability rates in the master equation[12]. Hence, the model for the constitutive gene becomes 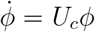, where *ϕ* = [*ϕ*_0_, *ϕ*_1_, …]^T^ and, e.g. [*U*_*c*_]_11_ = −*Nρ*, [*U*_*c*_]_12_ = −*ρ, U*_*c*_ is a ∞ × ∞ non-diagonal matrix, and its eigenvalues can be obtained indirectly from the equations governing the vector of moments *µ* of the stochastic process. Set *µ* = *Mϕ*. The entries of *M* are obtained from *µ* = [*µ*_0_, *µ*_1_,]^T^, where 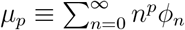 for *p* = 0, 1, …, and *µ*_0_ = 1. *µ* obeys the system of coupled ODEs 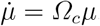. *Ω*_*c*_ is a lower triangular matrix where [*Ω*_*c*_]_*ii*_ = −(*i* − 1)*ρ, i* = 1, 2, Since *Ω*_*c*_ = *MU*_*c*_*M* ^−1^, *Ω*_*c*_ and *U*_*c*_ have the same characteristic polynomials −*λ*(−*λ* − *ρ*)(−*λ* − 2*ρ*), factorizable from *Ω*_*c*_ because of its shape. Thus *λ*_*l*_ = −*lρ, l* = 0, 1,, determine the temporal decay of the probabilities. It is hard to analytically obtain the roots of the characteristic polynomial of *U*_*c*_ because of its size and shape. That is done numerically truncating *U*_*c*_ size at some nite integer *K*. That will break the factorizability of the characteristic polynomial, and the first *L* (*L < K*) largest eigenvalues of the truncated *U*_*c*_ converge to the actual one (Fig. 1.A). This approach has limited power as we show below.

**Figure 1.**
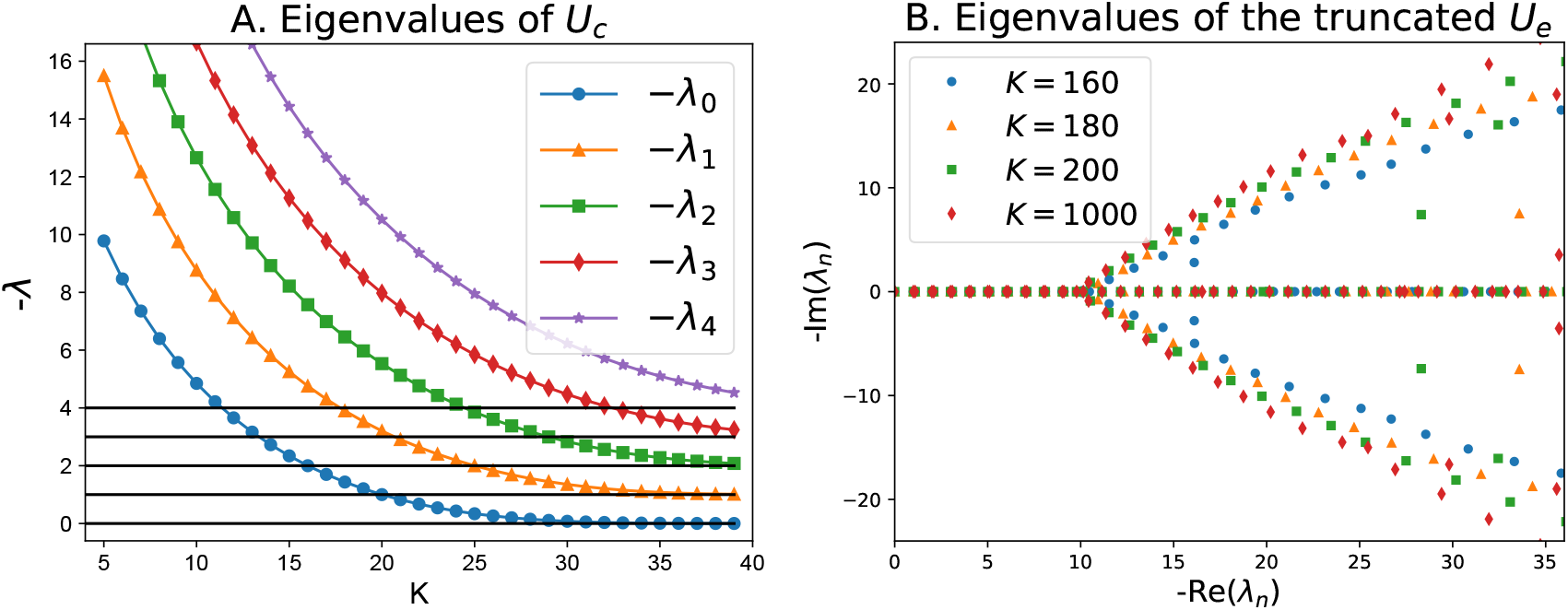
A. *k* = 20, *ρ* = 1. B. *k* = 20, *ρ* = 1, *f* = 4, *h* = 0.1.

Let us consider the master equation modeling the externally regulated (ERG) and self-repressing (SRG) gene[13]. The ON to OFF switching rate is *h* for the SRG and *h*_1_ for the ERG. *f* is the OFF to ON switching rate. *k* and *ρ* are, respectively, rates of protein synthesis when the promoter is ON and degradation. *α*_*n*_ and *β*_*n*_ obey:

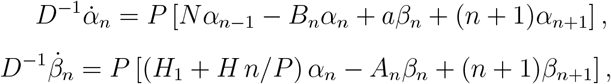

where 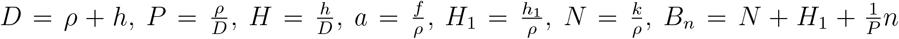, and *A*_*n*_ = *a* + *n*. For *D* = *ρ* and *P* = 1 we have an ERG and a SRG for *H*_1_ = 0.

Case 2: The marginal probability *ϕ*_*n*_ = *α*_*n*_+*β*_*n*_ replaces *β*_*n*_ such that *ψ* = [*ϕ*_0_, *α*_0_, *ϕ*_1_, *α*_1_, …]^T^. The master equation for the ERG becomes 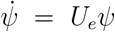, where *U*_*e*_ is the transition matrix. To compute the eigenvalues of *U*_*e*_ we take the moments vector 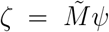, where *ζ* = [*µ*_0_, *ν* _0_, *µ*_1_, *ν* _1_, …]^T^. The entries of 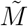 are built using 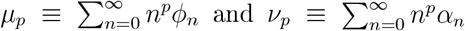. The moments vector obeys 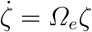, where *Ω*_*e*_ has lower triangular shape with [*Ω*_*e*_]_2*l*+1,2*l*+1_ = −*lρ* and [*Ω*_*e*_]_2*l*+2,2*l*+2_ = −(*ϵ* + *l*)*ρ* for *l* = 0, 1, Hence, the decaying rates of the stochastic process for the ERG are {−*lρ*} ∪{−(*ϵ* + *l*)*ρ*}, where *l* = 0, 1, … and 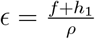. As suggested by the result of Case 1, one might attempt to numerically compute the roots of the characteristic polynomial of the truncated *U*_*e*_ instead of using the eigenvalues of *Ω*_*e*_. The characteristic polynomial of *Ω*_*e*_ and *U*_*e*_ factorizes as −*λ*(−*λ* − *ρ*)(−*λ* − *ϵ*) This factorizability is lost in the truncated matrix *U*_*e*_ and complex eigenvalues start to appear (Fig. 1.B). The eigenvalues of the truncated transition matrix agree with those obtained by continued fraction and that exempli es the methodological artifact underlying the complex eigenvalues reported in[10].

Case 3: the matrix *Ω*_*s*_ governing the moments vector of the SRG has a non-triangular shape. Thus we use its transition matrix *U*_*s*_. We de ne the diagonal matrix *Λ*, with *Λ*_2*l*,2*l*_ = −*lD* and *Λ*_2*l*+1,2*l*+1_ = −(*b* + *l*)*D*, where *l* = 0, 1, … and *b* = (*a* + *NH*)*P*. There is a matrix *V* obeying *V U*_*s*_ = *ΛV*, if the *Λ*_*ii*_’s are eigenvalues of *U*_*s*_. The rows of *V* are the left eigenvalues of *U*_*s*_ defined as a recurrence relation. *V*_*l*,0_ = *V*_0,*l*_ = 1, for *l* = 0, 1, For *i* = 1, 3, … we set: 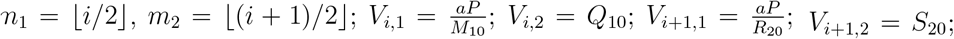 and for each *i*: *j* = 3, 5, … ; 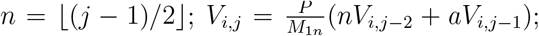 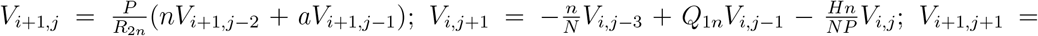 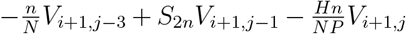; where 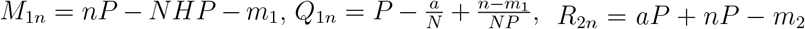, and 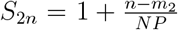. That is sufficient to demonstrate that the set of eigenvalues of *U*_*s*_ is {−*lD*} ∪ {−(*b* + *l*)*D*}, for *l* = 0, 1, Thus, our results are correct, and complete, as we computed the coefficients of the linear combination of concluent Heun functions for an initial condition as an inverse problem[1].

## IV. CONCLUDING REMARKS

The authors claimed to have found a new class of operators (their Eq. (6)) having complex eigenvalues and non-orthogonal eigenfunctions. That would lead to damped oscillating probability distributions which they do not show. This is because the imaginary decaying rates result from a methodological artifact. Indeed, it is well known that the operator of their Eq. (6), after multiplication by *w*(*z*) = *z*^*γ*^(*z* − 1)^*δ*^(*z* − *a*)^*ϵ*^, is written in the usual self-adjoint form *L*_*z*_*y* = *qw*(*z*)*y*[14].

## ACKNOWLEDGMENTS

AFR dedicates this document to Profs. JEM Hornos and J Reinitz, both in memorian, and thanks them, JCA Barata, M Rosner, G Balazsi, GCP Innocentini, AU Sabino, G Giovanini, B Carvalho, LR Gama for invaluable discussions. Research supported by Grant No. NIH R01 OD010936 and FUSP 403602.

